# ABCF ATPases involved in protein synthesis, ribosome assembly and antibiotic resistance: structural and functional diversification across the tree of life

**DOI:** 10.1101/220046

**Authors:** Victoriia Murina, Marje Kasari, Hiraku Takada, Mariliis Hinnu, Chayan Kumar Saha, James W. Grimshaw, Takahiro Seki, Michael Reith, Marta Putrinš, Tanel Tenson, Henrik Strahl, Vasili Hauryliuk, Gemma Catherine Atkinson

**Affiliations:** Department of Molecular Biology, Umeå University, 901 87, Umeå, Sweden; Laboratory for Molecular Infection Medicine Sweden (MIMS), Umeå University, 901 87, Umeå, Sweden; University of Tartu, Institute of Technology, Nooruse 1, 50411 Tartu, Estonia; Centre for Bacterial Cell Biology, Institute for Cell and Molecular Biosciences Newcastle University, Richardson Road, Newcastle upon Tyne, NE2 4AX, United Kingdom; Department of Applied Chemistry and Biotechnology, Faculty of Engineering, Chiba University, 263-8522, Chiba, Japan

## Abstract

Within the larger ABC superfamily of ATPases, ABCF family members eEF3 in *Saccharomyces cerevisiae* and EttA in *Escherichia coli* have been found to function as ribosomal translation factors. Several other ABCFs including biochemically characterised VgaA, LsaA and MsrE confer resistance to antibiotics that target the peptidyl transferase centre and exit tunnel of the ribosome. However, the diversity of ABCF subfamilies, the relationships among subfamilies and the evolution of antibiotic resistance factors from other ABCFs have not been explored. To address this, we analysed the presence of ABCFs and their domain architectures in 4505 genomes across the tree of life. We find 45 distinct subfamilies of ABCFs that are widespread across bacterial and eukaryotic phyla, suggesting they were present in the last common ancestor of both. Surprisingly, currently known antibiotic resistance (ARE) ABCFs are not confined to a distinct lineage of the ABCF family tree. This suggests that either antibiotic resistance is a pervasive feature of bacterial ABCFs, or it is relatively easy to evolve antibiotic resistance from other ABCF functions. Our data suggest there are a number of previously unidentified ARE ABCFs in antibiotic producers and important human pathogens. We also find that ATPase-deficient mutants of all four *E. coli* ABCFs (EttA, YbiT, YheS and Uup) inhibit protein synthesis, indicative of their ribosomal function, and demonstrate a genetic interaction of ABCFs Uup and YheS with translational GTPase BipA involved in assembly of the 50S ribosome subunit. Finally, we show that *Bacillus subtilis* VmlR is a ribosome-binding resistance factor localised to the cytoplasm.

**Author summary:** Isolated members of the ABCF protein family of ATP-hydrolysing enzymes have been found to have important roles in protein synthesis and antibiotic resistance. However, their full diversity across the tree of life, and their evolutionary histories have never been examined. Therefore, we analysed the presence of ABCFs and their constituent domains in genomes across the tree of life, discovering 45 distinct subfamilies of ABCFs that are widespread across bacterial and eukaryotic phyla. This includes several subfamilies that we predict comprise novel antibiotic resistance (ARE) ABCFs, present in antibiotic producers and important human pathogens. There are significant gaps in our knowledge about the functional capabilities of different ABCF families. To address this, we have made ATPase domain mutants of all four *Escherichia coli* ABCFs, showing that they inhibit protein synthesis and indicating a role on the ribosome. Furthermore, we demonstrate a genetic interaction of two *E. coli* ABCFs with the GTPase BipA, involved in ribosome assembly. Finally, we show that *Bacillus subtilis* VmlR in the ARE2 subfamily is a ribosome-binding resistance factor localised to the cytoplasm. As more is discovered about the function of individual ABCFs, the more it will be possible to predict functions of uncharacterised members, using the ABCF family tree as a framework.

## Introduction

Protein biosynthesis – translation – is the reading and deciphering of information coded in genes to produce proteins. It is one of the most ancient and central cellular processes, and control of the various stages of translation is achieved via an intricate interplay of multiple molecular interactions. For many years, enzymatic control of the ribosomal cycle was thought to be mainly orchestrated mainly by translational GTPases (trGTPases). That view of translation has been nuanced by the identification of multiple ATPases in the ABC superfamily that have important roles in translational regulation on the ribosome. The ABC protein eEF3 (eukaryotic Elongation Factor 3) is an essential factor for polypeptide elongation in *Saccharomyces cerevisiae* [1] with proposed roles in E-site tRNA release and ribosome recycling [2-4]. This fungi-specific translational ABC ATPase appeared to be an exception to the tenet that trGTPases are the enzymatic rulers of the ribosome, until ABCE1 (also known as Rli1), a highly conserved protein in eukaryotes and archaea, was identified as another ribosome recycling factor [5-7].

The ABC ATPases together comprise one of the most ancient superfamilies of proteins, evolving well before the last common ancestor of life [8]. The superfamily contains members with wide varieties of functions, but is best known for its membrane transporters [9]. Families of proteins within the ABC superfamily are named alphabetically ABCA to ABCH, following the nomenclature of the human proteins [10]. While most ABCs carry membrane-spanning domains (MSDs), these are lacking in ABCE and ABCF families [11]. ABCF proteins of eukaryotes include eEF3, and also other ribosome-associated proteins: Gcn20 is involved in sensing starvation by the presence of uncharged tRNAs on the eukaryotic ribosome [12]; ABC50 (ABCF1) promotes translation initiation in eukaryotes [13]; and both Arb1 (ABCF2) and New1 have been proposed to be involved in biogenesis of the eukaryotic ribosome [14, 15]. Ribosome-binding by ABCF proteins seemed to be limited to eukaryotes until characterisation of ABCF member EttA (energy-dependent translational throttle A) found in diverse bacteria. *Escherichia coli* EttA binds to the E-site of the ribosome where it is proposed to ‘throttle’ the elongation stage of translation in response to change in the intercellular ATP/ADP ratio [16, 17]. EttA is one of four *E. coli* ABCFs, the others being YheS, Uup and YbiT, none of which have yet been shown to operate on the ribosome [16].

Bacterial ABCF family members have been found to confer resistance to ribosome-inhibiting antibiotics widely used in clinical practise, such as ketolides [18], lincosamides [19-22], macrolides [23, 24], oxazolidinones [25], phenicols [25], pleuromutilins [22] and streptogramins A [22, 26] and B [24]. These <u>a</u>ntibiotic <u>re</u>sistance (AREs) ABCFs have been identified in antibiotic-resistant clinical isolates of *Staphylococcus, Streptomyces* and *Enterococcus* among others [27]. This includes the so-called ESKAPE pathogens *Enterococcus faecium* and *Staphylococcus aureus* that contribute to a substantial proportion of hospital-acquired multidrug-resistant infections [28]. As some efflux pumps carry the ABC ATPase domain, it was originally thought that ARE ABCFs similarly confer resistance by expelling antibiotics. However, as they do not carry the necessary transmembrane domains, this is unlikely [29, 30]. In support of this, it was recently shown that *Staphylococcus aureus* ARE VgaA protects protein synthesis activity in cell lysates from antibiotic inhibition and that *Enterococcus faecalis* ARE LsaA displaces radioactive lincomycin from *S. aureus* ribosomes [31]. Using a reconstituted biochemical system, we have shown that VgaA and LsaA directly protect the ribosome peptidyl transferase centre (PTC) from antibiotics in an ATP-dependent manner [32]. Recent cryo-EM structures of AREs *Pseudomonas aeruginosa* MsrE and *Bacillus subtilis* VmlR on the ribosome show that like EttA, these ABCFs bind to the E-site of the ribosome, with extended inter-ABC domain linkers protruding into the PTC [33, 34]. The ARE ABCFs therefore appear to either physically interact with the drug to displace it from the ribosome, or allosterically induce a change in conformation of the ribosome that ultimately leads to drug drop-off [35-37].

Here, we carry out an in-depth survey of the diversity of ABCFs across all species with sequenced genomes. We find 45 groups (15 in eukaryotes and 30 in bacteria), including 7 groups of AREs. So-called EQ_2_ mutations, double glutamic acid to glutamine substitutions in the two ATPase active sites of EttA lock the enzyme on ribosome in an ATP-bound conformation, inhibiting protein synthesis and cellular growth [16, 17]. We have tested the effect of equivalent mutations in the other three *E. coli* ABCFs – YbiT, YheS and Uup – as well as the *Bacillus subtilis* ARE VmlR. We establish genetic associations of *E. coli* ABCFs YheS and Uup with the translational GTPase BipA (also known as TypA), and through microscopy and polysome profile analyses, confirm that VmlR does not confer lincomycin resistance through acting as a membrane-bound pump, but via direct interaction with cytoplasmic ribosomes.

## Results

### ABCFs are widespread among bacteria and eukaryotes

To identify candidate subfamilies of ABCFs and refine the classifications an iterative bioinformatic protocol of sequence searching and phylogenetic analysis was applied. Sequence searching was carried out against a local database of 4505 genomes from across the tree of life. To first get an overview of the breadth of diversity of ABCFs across life, sequence searching began with a local BlastP search against a translated coding sequence database limited by taxonomy to one representative per class, or order if there was no information on class for that species in the NCBI taxonomy database. From phylogenetic analysis of the hits, preliminary groups were identified and extracted to make Hidden Markov Models (HMMs) for further sequence searching and classification. Additional sequences from known ABCF AREs were included in phylogenetic analyses in order to identify groups of ARE-like ABCFs. HMM searching was carried out at the genus level followed by phylogenetic analysis to refine subfamily identification, with final predictions made at the species level. The resulting classification of 16848 homologous sequences comprises 45 subfamilies, 15 in eukaryotes and 30 in bacteria. Phylogenetic analysis of representatives across the diversity of ABCFs shows a roughly bipartite structure, with most eukaryotic sequences being excluded from those of bacteria with strong support (fig. 1). Five eukaryotic groups that fall in the bacterial part of the tree are likely to be endosymbiotic in origin (see the section *Bacteria-like eukaryotic ABCFs*, below).

**Fig. 1.**
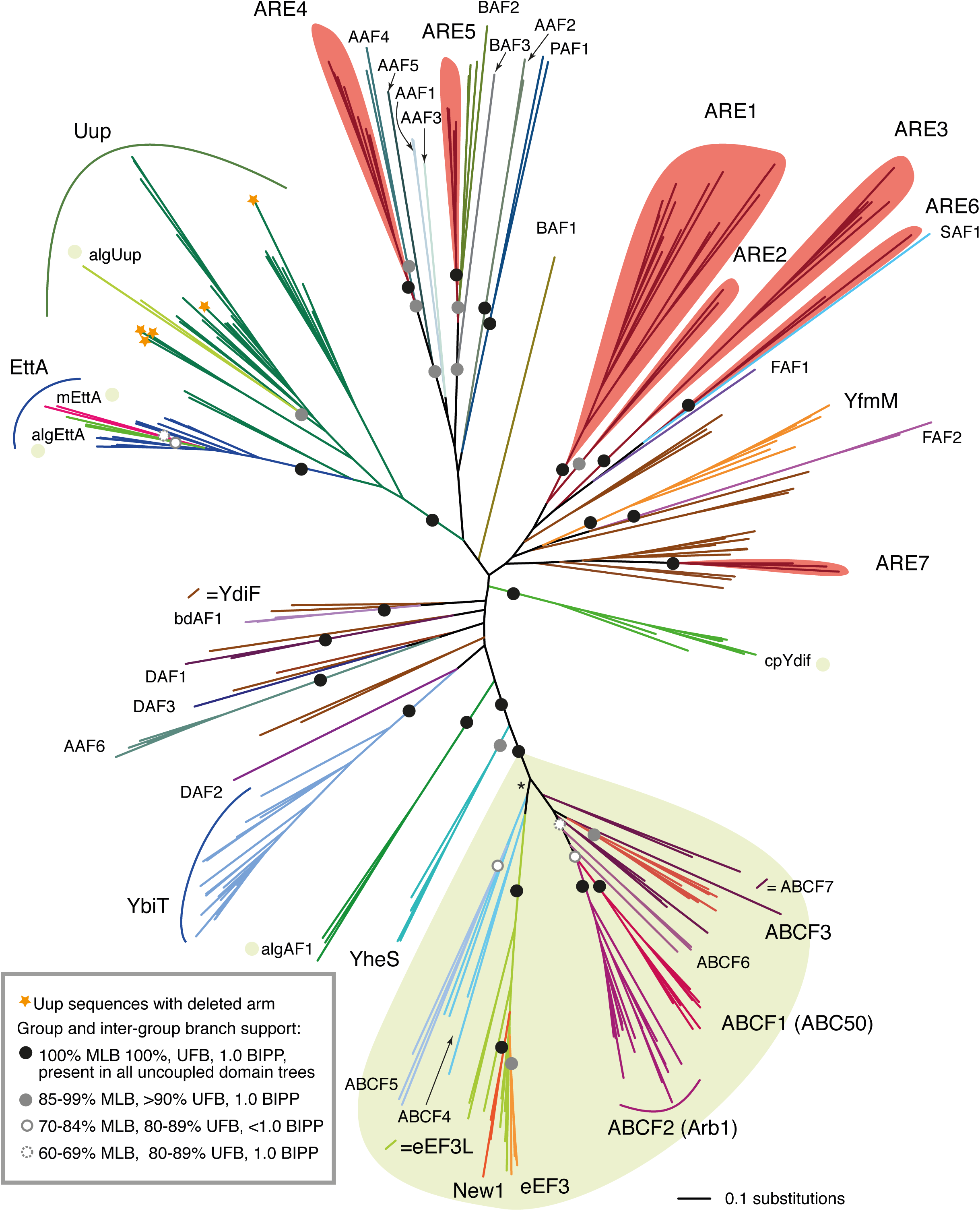
The family tree of ABCFs has a bipartite structure corresponding to eukaryotic-like and bacterial (and organellar)-like sequences. The tree is a RaxML maximum likelihood phylogeny of representatives across the ABCF family with branch support values from 100 bootstrap replicates with RaxML (MLB), 1000 ultrafast bootstrap replicates with IQ-TREE (UFB) and Bayesian inference posterior probability (BIPP). The inset box shows the legend for subfamily and intersubfamily support; support values within subfamilies and that are less that 60% MLB are not shown. Species were chosen that sample broadly across the tree of ABCF-encoding life, sampling at least one representative from each subfamily. Green shading shows the eukaryotic type ABCFs; other subgroups are bacterial unless marked with a green shaded circle to indicate eukaryotic groups with potentially endosymbiotic origin. CpYdif contains both cyanobacterial and predicted chloroplast sequences. The full tree with taxon names and sequence IDs is shown in fig. S1. Branch lengths are proportional to amino acid substitutions as per the scale bar in the lower right. The asterisked branch is not supported by this data set, however it is supported at 85% MLB in phylogenetic analysis of the eukaryotic subgroup and its viral relatives, rooted with YheS (fig. S3). Branch lengths are proportional to amino acid substitutions as per the lower right scale bar.

ABCFs are widespread among bacteria and eukaryotes; there are on average four ABCFs per bacterial genome, and five per eukaryotic genome. However, there is considerable variation in how widespread each subfamily is (table 1). The presence of all subfamilies in each genome considered here is shown in table S1, with the full set of sequence IDs and domain composition recorded in table S2. Domain coordinates by amino acid position can be found in table S3. In bacteria, Actinobacteria, and Firmicutes are the phyla with the largest numbers of ABCFs (up to 11 per genome), due to expansions in ARE and potential novel ARE ABCF subfamilies. Among eukaryotes plants and algae encode the most subfamilies, probably due in part to gene acquisition from endosymbiosis events. The diatom *Fragilariopsis cylindrus* has 30 ABCFs and the Haptophyte *Emiliania huxleyi* has 26. Bacterial contamination can sometimes inflate the number of genes in eukaryotic genomes as noted previously for trGTPases [35]. However, as all the *Fragilariopsis* and *Emiliania* sequences belong to typically eukaryotic subgroups, they do not appear to be the result of bacterial contamination. The Tibetan antelope *Pantholops hodgsonii*, on the other hand has 25 ABCFs, 20 of which belong to bacterial subgroups. The genome is known to be contaminated by *Bradyrhizobium* sequences [38], and thus the bacteria-like hits from *P. hodgsonii* are most likely artifacts.

**Table 1.**
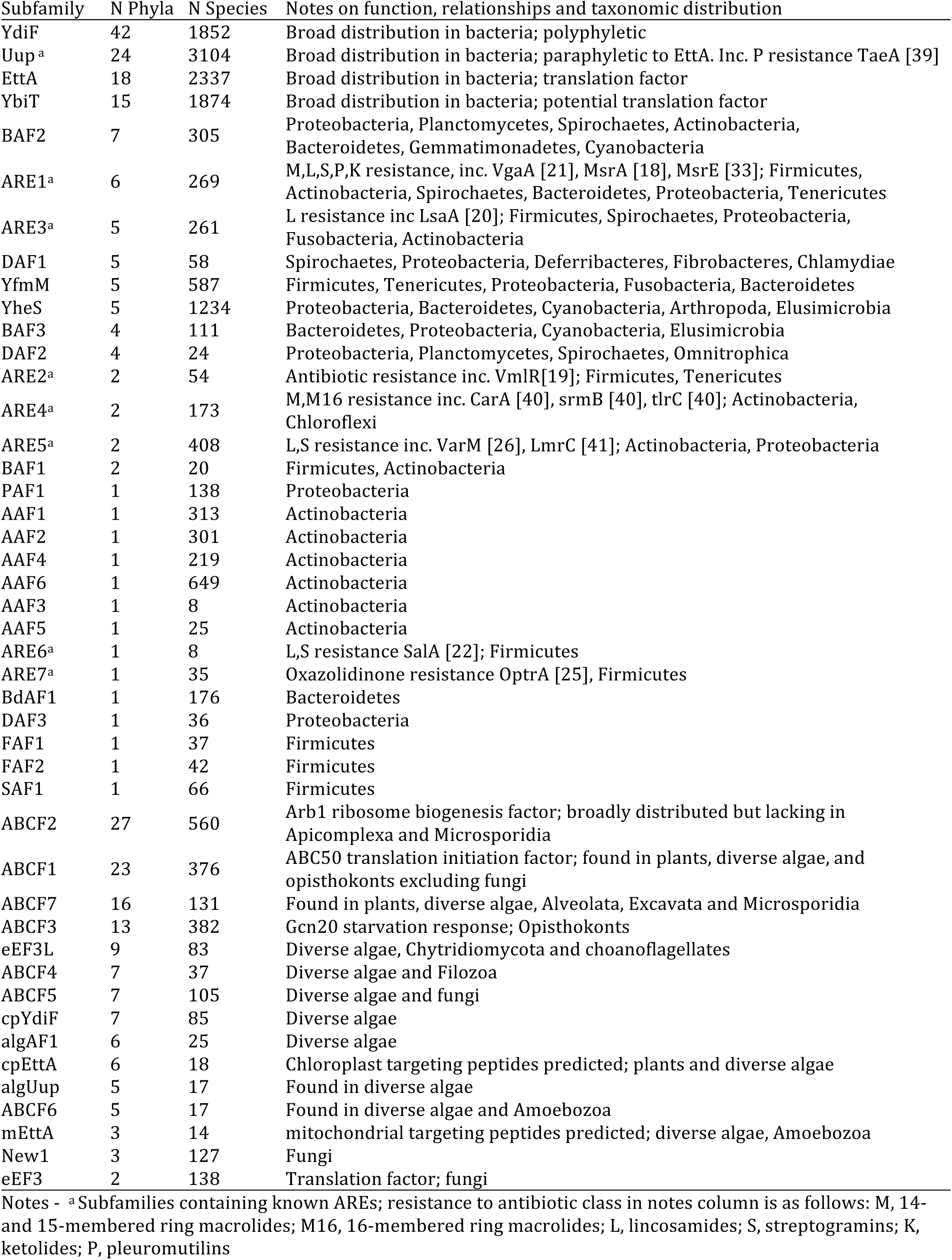
The subfamilies of the ABCF family, and the numbers (N) of phyla and species in which they are encoded.

The wide distribution and multi-copy nature of ABCFs suggests an importance of these proteins. However they are not completely universal, and are absent in almost all Archaea. The euryarcheaotes *Candidatus Methanomassiliicoccus intestinalis, Methanomethylophilus alvus, Methanomassiliicoccus luminyensis*, and *Thermoplasmatales archaeon BRNA1* are the only archaea found to encode ABCFs, in each case YdiF. ABCFs are also lacking in 214 bacterial species from various phyla, including many endosymbionts. However, the only phylum that is totally lacking ABCFs is Aquificae. ABCFs are almost universal in Eukaryotes; the only genomes where they were not detected were those of Basidomycete *Postia placenta*, Microsporidium *Enterocytozoon bieneusi*, and apicomplexan genera *Theileria, Babesia* and *Eimeria*.

Many of the phylogenetic relationships among subfamilies of ABCFs are poorly resolved (fig. 1). This is not surprising, since these represent very deep bacterial relationships that predate the diversification of major phyla and include gene duplication, and differential diversification and loss, and likely combine both vertical inheritance and horizontal gene transfer. Although relationships can not be resolved among all subfamilies, some deep relationships do have strong support (i.e. maximum likelihood bootstrap percentage (MLB) of more than 85% and Bayesian inference posterior probability (BIPP) of 1.0); EttA and Uup share a common ancestor to the exclusion of other ABCFs with full support (100% MLB, 100% UFB and 1.0 BIPP fig. 1, fig. S1) and YheS is the closest bacterial group to the eukaryotic ABCFs with strong support (94% MLB, 98% UFB, and 1.0 BIPP; (fig. 1, fig. S1, text S1). This latter observation suggests that eukaryotic-like ABCFs evolved from within the diversity of bacterial ABCFs. However, this depends on the root of the ABCF family tree. To address this, phylogenetic analysis was carried out of all ABCFs from *E. coli, Homo sapiens, S. cerevisiae* and *B. subtilis*, along with ABCE family sequences from the UniProt database [42]. Rooting with ABCE does not provide statistical support for a particular group at the base of the tree, but does support the eukaryotic subgroups being nested within bacteria, with YheS as the closest bacterial group to the eukaryotic types (fig. 2). It also shows that eEF3 and New1 are nested within the rest of the ABCF family, thus confirming their identity as ABCFs, despite their unusual domain structure. To address the possibility that due to recombination the two ABC domains of the ABCF family may have had different evolutionary histories, we repeated our phylogenetic analysis of representative sequences with the ABC domains uncoupled and aligned to each other (fig. 1, Text S1). With this very short alignment (204 positions) containing a larger proportion of almost invariant active site residues, there is even less statistical support for relationships among subgroups. Nevertheless, we still retain the branches that are well supported in our full-length analyses (fig. 1), and thus there is no evidence for recombination.

**Fig. 2.**
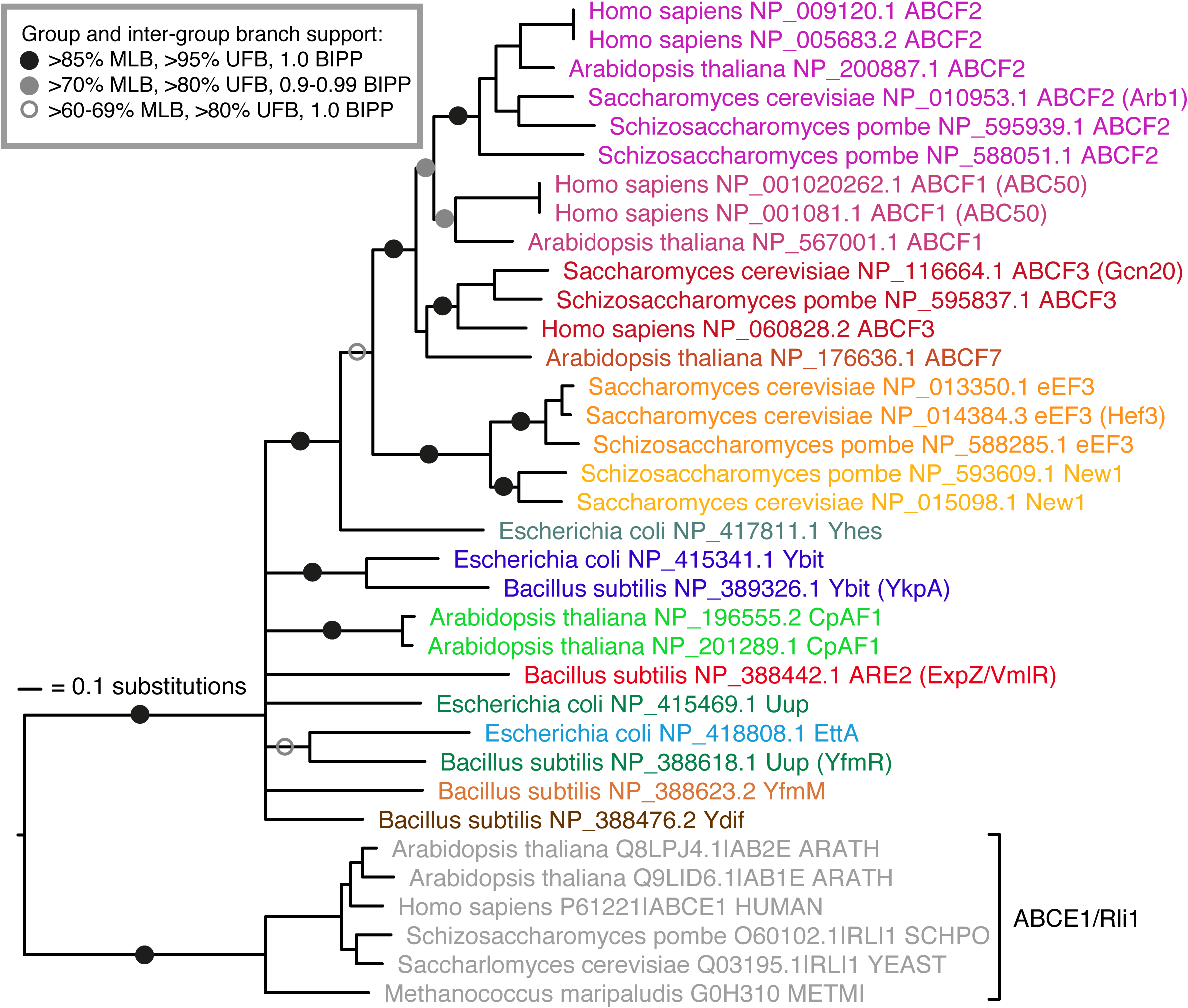
Rooting with ABCE shows eukaryotic-like ABCFs nesting within bacterial-like ABCFs, with YheS as the sister group to the eukaryotic-like clade. Maximum likelihood phylogeny of representatives across the ABCF family, and ABCE sequences from the UniProt database. Branch support from 200 bootstrap replicates with RaxML (MBP), 1000 ultrafast bootstrap replicates with IQ-TREE and Bayesian inference posterior probability is indicated with the key in the inset box. Branch lengths are proportional to amino acid substitutions as per the inset scale bar.

### Domain architectures in the ABCF family are variable

To assess the conservation of domains across the family tree of ABCFs, we extracted the domain regions from subfamily alignments, made HMMs representing each domain region and scanned every sequence in our database. We find the most common domain and subdomain structure is an N-terminal ABC1 nucleotide binding domain (NBD) containing an internal Arm subdomain, followed by the Linker region joining to the ABC2 NBD (fig. 3A). Variations on this basic structure include deletions in the Arm and Linker regions, insertion of a Chromo subdomain in the ABC2 NBD, and extension of N and C termini by sequence extensions (fig. 3A). The domain structures of eukaryotic ABCFs are more diverse than those of bacteria, with greater capacity for extensions of the N-terminal regions to create new domains (fig. 3A). An increased propensity to evolve extensions, especially at the N terminus is also seen with eukaryotic members of the trGTPase family [35]. In bacteria, terminal extensions of ABCFs tend to be at the C terminus (fig. 3A-B).

**Fig. 3.**
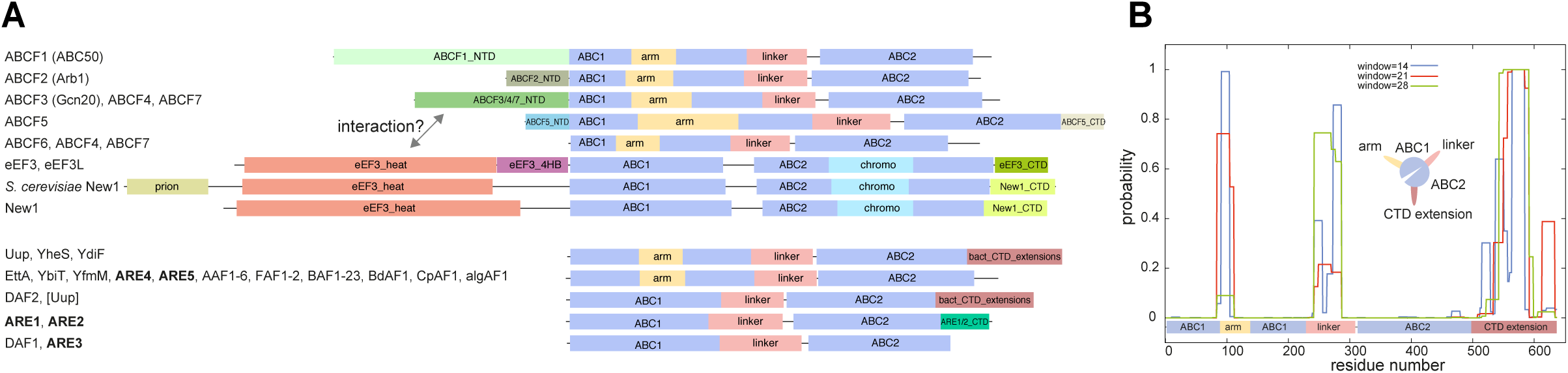
Typical domain and subdomain architectures of ABCFs. (A) Boxes show domains as predicted by HMMs. Full coordinates and sequence data for these examples are recorded in Table S3. Dotted lines shows possible interactions between the HEAT domain of eEF3/New1/eEF3L and the N-terminal domain of ABCF3, ABCF4 and ABCF7. (B) Predicted coiled coil regions of *E. coli* YheS along the protein length. Inset: cartoon representation of the coiled coil subdomains protruding from the core ABC domains.

Cryo-electron microscopy structures show that bacterial ABCFs bind to the same site of the ribosome, which is different from that of eEF3 [2, 17, 33, 34], and this is reflected in their domain architectures. eEF3 carries additional N-terminal domains and does not have the Arm subdomain that in EttA binds the L1 stalk and ribosomal protein L1. Instead it carries a Chromo (Chromatin Organization Modifier) subdomain in ABC2 that – from a different orientation to the Arm domain of EttA - interacts with, and stabilises the conformation of the L1 stalk [2]. The Arm subdomain is also missing in eEF3-like close relatives New1 and eEF3L. Although the Arm is widespread in bacterial ABCFs, it is not universal; it is greatly reduced in a number of subfamilies, with the most drastic loss seen in ARE1 and 2 (figs. 3A, and see *Putative AREs* section, below). Subdomain composition can even vary within subfamilies; Uup has lost its arm independently in multiple lineages (fig. 1, fig. S1).

Arms, linkers and CTD extensions are poorly conserved at the primary sequence level, but are similar in terms of composition, all being rich in charged amino acids, particularly arginine and lysine. Their variable presence and length suggests they can be readily extended or reduced during evolution. The CTD of Uup forms a coiled coil structure that is capable of binding DNA [43]. However, whether DNA binding is its primary function is unclear. The CTD of YheS has significant sequence similarity (E value 3.58e-03) to the tRNA binding CTD of Valine-tRNA synthetase (NCBI conserved domains database accession cl11104). *In silico* coiled coil prediction suggests there is a propensity of all these regions to form coiled coil structures (fig. 3B). Extensions and truncations of the arms, linkers and CTD extensions possibly modulate the length of coiled coil protrusions that extend from the globular mass of the protein.

### Eukaryotic ABCFs comprise 15 subfamilies

#### eEF3, New1 and eEF3L

eEF3, eEF3L and New1 group together and are particularly distinct members of the ABCF family. The eEF3 subfamily represents the classical fungal proteins, while eEF3L is a more divergent group found mainly in protists (see below). eEF3 has a recent paralogue in *S. cerevisiae* (Hef3 (fig. 2), YEF3B) that apparently arose as a result of the whole genome duplication in yeast [44, 45]. The conservation of domain structure in eEF3/New1/eEF3L suggests they bind the ribosome similarly (fig. 3A). Ribosome binding by eEF3 involves the Chromo subdomain, and the HEAT domain [2], which in addition to New1 and eEF3L, is also found in the protein Gcn1, a binding partner of Gcn20 ABCF [46] (see the section *ABCF1-7*, below). eEF3 has a distinct C-terminal extension (fig. 3A), through which it interacts with eEF1A [47]. It has been suggested that this leads to the recruitment of eEF1A to *S. cerevisiae* ribosomes [47, 48]. The New1 CTD contains a region of sequence similarity to the eEF3 CTD; both contain a polylysine/arginine-rich tract of 20-25 amino acids (fig. S2). In *S. cerevisiae* eEF3, this is at positions 1009-1031, which falls within the eEF1A-binding site. Thus eEF1A binding may be a common feature of eEF3, New1, and also eEF3L, which commonly includes an eEF3-like C-terminal extension (table S2).

We find eEF3-like (eEF3L) factors in a range of eukaryotes including choanoflagellates, haptophytes, heterokonts, dinoflagellates, cryptophytes and red and green algae. This suggests the progenitor of eEF3 was an ancient protein within eukaryotes, and has been lost in a number of taxonomic lineages. Alternatively, eEF3L may have been horizontally transferred in eukaryotes. eEF3 is found in *Chlorella* viruses [49] and *Phaeocystis* viruses (fig. S3), suggesting this may be a medium of transfer. There is no “smoking gun” for viral-mediated transfer in the phylogenies, in that eukaryotic eEF3L sequences do not nest within viral sequence clades (fig. S3). However, given the close association of viral and protist eEF3L, this still remains a possibility. Curiously, the taxonomic distribution of eEF3L in diverse and distantly related protists is similar to that of the unusual elongation factor 1 (eEF1A) paralogue EFL [50] (table S4). The propensity for eEF3/eEF3L/New1 to be present in EFL-encoding organisms and absent in eEF1A-encoding organisms is significant at the level of *P*<0.0001 with Fisher’s exact test. Like eEF3L, EFL can be found in viruses such as *Aureococcus anophagefferens* virus (NCBI protein accession YP_009052194.1). As eEF3 interacts with eEF1A [48], the equivalents eEF3L and EFL may also interact in the organisms that encode them.

Like eEF3, New1 is found across the fungal tree of life (table S1, fig. S3). *S. cerevisiae* New1 has previously been reported to carry a prion-like Y/N/Q/G repetitive region in the N-terminal region before the HEAT domain [51]. We find that this region is limited in taxonomic distribution to Saccharomycetale yeast (table S2, fig. 3A and fig. S2). Thus it is not found in the N-terminal region of *Schizosaccharomyces pombe* New1 (also known as Elf1).

#### ABCF1-7

ABCF1-7 comprise the “ancestral-type” eukaryotic ABCFs, in that they have the typical ABC domain structure that is seen in bacterial ABCFs, and they lack the Chromo and HEAT domains that are found in eEF3, New1 and eEF3L (fig. 3A). All of the terminal extensions found in ABCF subfamilies are biased towards charged amino acids, often present as repeated motifs. The ABCF1 (ABC50) NTD (which interacts with eIF2 [52]) is, due to its length and number of repeats, one of the most striking (fig. S2). It contains multiple tracts of poly-lysine/arginine, poly–glutamic acid/aspartic acid and – in animals - poly-glutamine. ABCF1 and ABCF2 have moderate support as sister groups (fig. 1), and both have representatives in all eukaryotic superphyla, but are not universal. Notable absences of ABCF1 are fungi, amoebozoa and most Aves (birds) (table 1, table S1).

ABCF2 (Arb1) is essential in yeast and its disruption leads to abnormal ribosome assembly [14], and human ABCF2 can complement an Arb1 deletion, suggesting conservation of function [53]. The protein is broadly distributed across eukaryotes, with the notable exception of the *Alveolata* superphylum (Table S1). Like other ABCF terminal extensions, the ABCF2 N-terminal domain is rich in lysine, in this case lysine and alanine repeats.

In yeast, where it is known as Gcn20, ABCF3 is a component of the general amino acid control (GAAC) response to amino acid starvation, acting in a complex with Gcn1 [54]. Gcn20 binds to Gcn1 via the latter’s HEAT-containing N-terminal domain [46], an interaction that is conserved in ABCF3 and Gcn1 of *Caenorhabditis elegans* [55]. As the HEAT domain is also found in eEF3, this raises the possibility that Gcn20/ABCF3 and eEF3 interact in encoding organisms (fig. 3A). Possible support for this comes from the observation that eEF3 overexpression impairs Gcn2 activation [56].

ABCF3 is widespread in eukaryotes, but absent in heterkont algae and archaeplastida. However, these taxa encode ABCF4 and ABCF7 of unknown function, which have N-terminal domains homologous to ABCF3. Thus, ABCF4 and ABCF7 may be the functional equivalents of ABCF3 in these taxa, potentially interacting with HEAT domain-containing eEF3L in organisms that encode the latter (fig. 3A). ABCF3 from four fungi (*Setosphaeria turcica,* NCBI protein accession number XP_008030281.1; *Cochliobolus sativus,* XP_007703000.1; *Bipolaris oryzae,* XP_007692076.1; and *Pyrenophora teres,* XP_003306113.1) are fused to a protein with sequence similarity to WHI2, an activator of the yeast general stress response [57].

ABCF5 is a monophyletic group limited to fungi and green algae (*Volvox* and *Chlamydomonas*), with a specific NTD and CTD (fig. 1 and 2A). ABCF5 is found in a variety of Ascomycete and Basidiomycete fungi, including the yeast *Debaryomyces hansenii*, but is absent in yeasts *S. pombe* and *S. cerevisiae*. ABCF4 is a polyphyletic group of various algal and amoeba protists that can not be assigned to the ABCF5, or eEF3/eEF3L/New1 clades. In eukaryotic ABCF-specific phylogenetic analysis, ABCF4/ABCF5/eEF3/eEF3L/New1 are separated from all other eukaryotic ABCFs with moderate support (85% MLB fig. S3). ABCF6 represents a collection of algal and amoebal sequences that associate with the ABCF1+ABCF2 clade with mixed support in phylogenetic analysis (fig. 1).

#### Bacterial-like eukaryotic ABCFs

Five eukaryotic subfamilies are found in the bacteria-like subtree: algAF1, algUup, mEttA, algEttA and cpYdiF (fig. 1). Given their affiliation with bacterial groups, they may have entered the cell with an endosymbiotic ancestor. Indeed, chloroplast-targeting peptides are predicted at the N termini of the majority of cpYdiF sequences, and mitochondrial localization peptides at most mEttA N termini. The situation is less clear for the three remaining groups, with a mix of signal peptides, or none at all being predicted across the group members (table S5).

### Bacterial ABCFs comprise 30 subfamilies, most of which have unknown function

There are 30 groups of bacterial ABCFs, the most broadly distributed being YdiF (the subfamily is given the name of the *Bacillus subtilis* protein as it is not present in *E. coli*) (table 1). This subfamily is a paraphyletic grouping comprising ABCF sequences that can not confidently be classified into any of the other subgroups (fig. 1). The next most broadly distributed group is Uup (*B. subtilis* protein name YfmR), which itself is paraphyletic to EttA. *B. subtilis* does not encode EttA, but does encode YbiT (*B. subtilis* name YkpA). It also encodes two ABCFs not present in *E. coli*: YfmM and VmlR (also known as ExpZ). VmlR is in the ARE2 subfamily, and confers resistance to virginiamycin M1 and lincomycin [19]. Insertional disruptants of all the chromosomal ABCF genes in *B subtilis* strain 168 have been examined for resistance to a panel of nine MLS class antibiotics, and only VmlR showed any hypersensitivity [19]. With the exception of EttA [16, 17], and the seven antibiotic resistance ARE ABCFs (table 1), the biological roles of the other 22 bacterial ABCFs are largely obscure.

#### B. subtilis *ARE VmlR is a cytoplasmic protein that directly protects the ribosome from antibiotics*

*B. subtilis* virginiamycin M and lincomycin resistance factor ABCF VmlR was originally annotated as an ABC efflux transporter, i.e. a membrane protein [19, 58]. To probe VmlR’s interaction with the ribosome we took advantage of ATPase-deficient VmlR mutants generated by simultaneous mutation of both glutamate residues for glutamine (EQ_2_) [59] that lock ABC enzymes in an ATP-bound active conformation [17, 60]. In the case of EttA, expression of the EQ_2_ mutant results in a dominant-negative phenotype as EttA incapable of ATP hydrolysis acts as a potent inhibitor of protein synthesis and, consequently, bacterial growth [16]. We constructed C-terminally tagged His_6_-TEV-3xFLAG-tagged (HTF-tagged) wild type and EQ_2_ (*vmlR*-HTF and *vmlR*EQ_2_-HTF) under the control of an IPTG-inducible P_*hy-spank*_ promotor [61]. To probe the intracellular localization of VmlR, we C-terminally tagged VmlR with the mNeonGreen fluorescent protein [62] under the control of xylose inducible promoter protein P_xyl_ [63]. We have validated the functionality of the fusion constructs by lincomycin resistance assays using a Δ*vmlR* knock-out strain as a negative control and a Δ*vmlR* knock-out strain expressing untagged VmlR under the control of an IPTG-inducible P_*hy-spank*_ promotor as a positive control (fig. S4A-C). While C-terminal tagging with either HTF or mNeonGreen does not abolish VmlR’s activity, the EQ_2_ versions of the tagged proteins are unable to protect from lincomycin (fig. S4B).

After establishing the functionality of the tagged VmlR constructs, we tested the effects of expression of either wild type or EQ_2_ VmlR-HTF on *B. subtilis* growth in rich LB media (fig. 4A). Expression of the wild type protein in the Δ*vmlR* background has no detectable effect. In contrast, the EQ_2_ version inhibits growth: while exponential growth is unaffected, the cells enter the stationary phase at lower cell densities, abruptly stopping growth instead of slowing down gradually (fig. 4A). Two factors are likely to cause the growth-phase specificity of the inhibitory effect. First, during the exponential growth cells efficiently dilute the toxic protein via cell division, and when the growth slows down, VmlR-EQ_2_-HTF accumulates. Second, upon entering the early stationary phase, *B. subtilis* sequesters 70S ribosomes into inactive 100S dimers [64], and this decrease of active ribosome concentration could conceivably render the cells more vulnerable to the inhibitory effects of VmlR-EQ_2_. We probed the interaction of wild type and EQ_2_ VmlR-HTF with ribosomes using polysome analysis in sucrose gradients in combination with Western blotting (fig. 4B). While the wild type protein barely enters the gradient (most likely dissociating from the ribosomes during centrifugation), the EQ_2_ version almost exclusively co-localises with the 70S peak fraction and is absent from the polysomal fractions (fig. 4B), suggesting co-sedimentation of a tight 70S:VmlR-EQ_2_ complex.

**Fig. 4.**
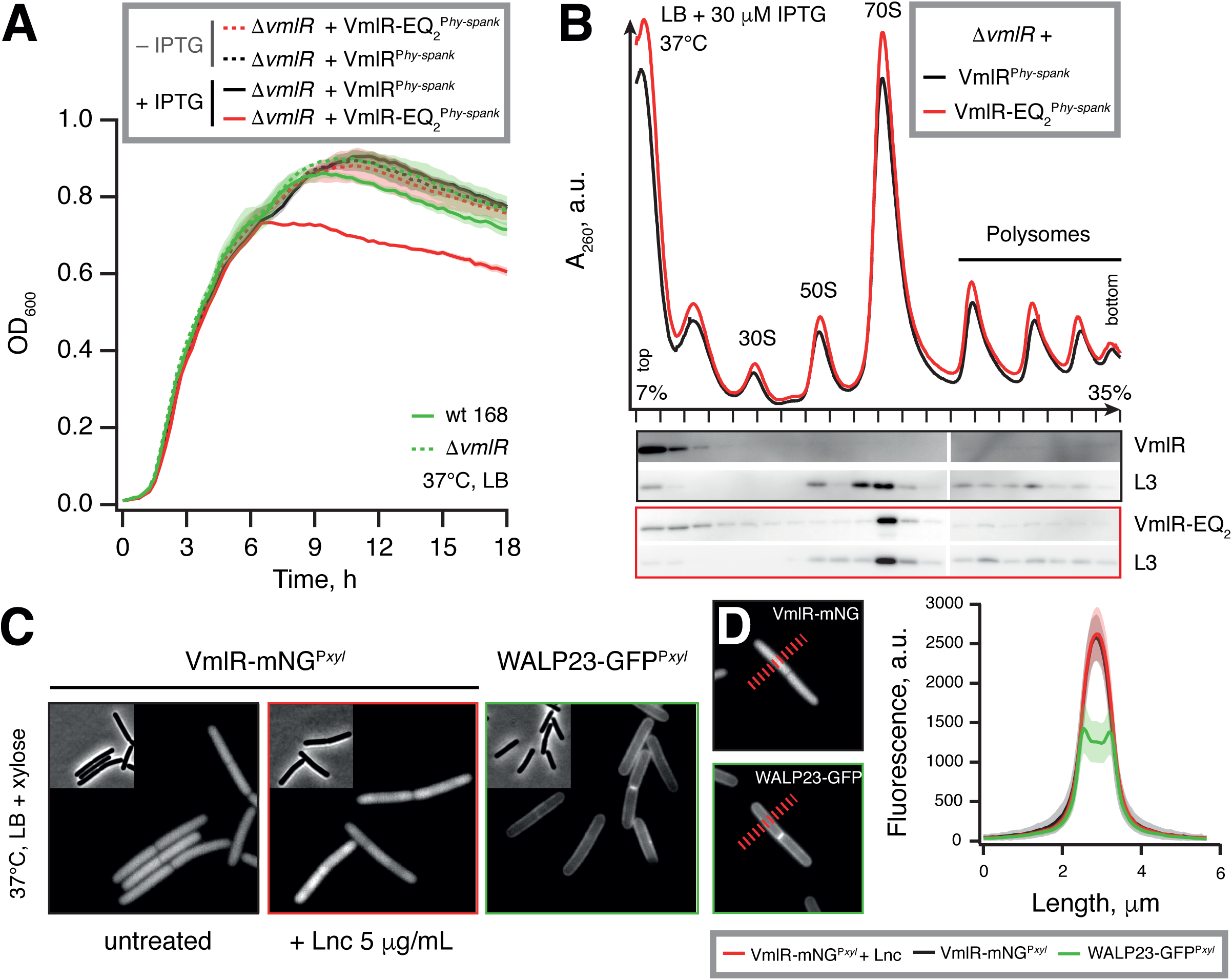
*B. subtilis* ARE VmlR is a cytoplasmic protein that directly protects the ribosome from antibiotics. (A) Growth of wild type *B. subtilis* 168, isogenic Δ*vmlR* knockout as well as Δ*vmlR* knockout expressing either wild type or EQ_2_ version of VmlR under the control of IPTG-inducible P_*hy-spank*_ promoter. Six biological replicates were averaged for each growth curve and the data presented as geometric means ± standard deviation. (B) Polysome analysis and western blotting of Δ*vmlR B. subtilis* expressing C-terminally HTF-tagged wild type and EQ_2_ version of VmlR. (C) Phase contrast and fluorescence images of uninhibited *B. subtilis* cells expressing VmlR-mNeonGreen (VmlR-mNG) in the presence and absence of lincomycin (40 min incubation with 5 µg/ml), and a model transmembrane protein WALP23-GFP are shown for comparison. (D) Fluorescence intensity profiles were measured perpendicular to the cell length axis along a 325 nm wide and 5.8 µm long line as indicated. Fluorescence intensity profiles of cells expressing WALP23-GFP [65], and cells expressing VmlR-mNG in the presence and absence of lincomycin. The graph depicts the average fluorescence intensity profiles and the corresponding standard deviations (n = 30).

Finally, having ascertained the functionality of VmlR-mNeonGreen (fig. S4C), we imaged *B. subtilis* cells expressing VmlR-mNeonGreen in the presence and absence of lincomycin (fig. 4C) and quantified the intensity of the fluorescent signal across the cell (fig. 4D). As a positive control for membrane localization we used WALP23-GFP [65] – an artificial model transmembrane helix WALP23 [66] fused with an N-terminal GFP label. We observe no evidence for association of VmlR with the membrane: the protein is clearly cytoplasmic, with a slight exclusion from the nucleoid in the presence of 5 μg/mL lincomycin. A likely explanation for this effect is the nucleoid compaction caused by inhibition of translation resulting in protein exclusion from the nucleoid-occupied space [67-69]. However, we cannot rule out that this general effect potentiated by specific interaction of VmlR with strongly nucleoid-excluded ribosomes.

#### E. coli *ABCFs EttA, YbiT, YheS and Uup interact genetically and functionally with protein synthesis and ribosome assembly*

*E. coli* encodes four ABCFs: EttA, YbiT, YheS and Uup. An array of structural, biochemical and microbiological methods has been used to establish that EttA operates on the ribosome [16, 17]. Ribosomal association of the other *E. coli* ABCFs has not been shown. However, a recent PhD thesis by Dr. Katharyn L. Cochrane suggests that Uup genetically interacts with an enigmatic ribosome-associated factor, the translational GTPase BipA (TypA) [70]. The *E. coli bipA* knock-out strain is characterised by a decreased level of 50S subunits accompanied by an accumulation of pre-50S particles [71]. In the presence of its native substrate, GTP, BipA associates with mature 70S ribosomes [72], occupying the ribosomal A-site [73]. However, in the presence of stress alarmone (p)ppGpp – a molecular mediator of the stringent response [74] – BipA binds the 30S subunit [75].

We have set out to systematically probe the involvement of *E. coli* ABCFs in protein synthesis. We used two experimental systems. The first is geared towards low level constitutive expression of native, untagged wild type and EQ_2_ proteins in a clinically relevant uropathogenic *E. coli* strain CFT073 [76]. For this, we cloned ABCF genes into a low copy pSC101 vector under control of a constitutive tet-promoter (P_tet_) that in the original plasmid drives expression of the tetracycline efflux pump TcR [77]. Using the λRed-mediated gene disruption method [78] we generated a set of mutants of lacking each of the four ABCF genes, as well as a Δ*bipA* and Δ*bipA*Δ*uup* knock-out strains. The second system allows inducible high-level expression of tagged proteins in the avirulent BW25113 *E. coli* strain [79, 80]. We used wild type and EQ_2_ mutants of EttA, YbiT, YheS and Uup with N-terminal FLAG-TEV-His_6_ (FTH)-tags expressed from a low copy pBAD18 plasmid under an arabinose-inducible araBAD (P_BAD_) promoter. The BW25113 strain can not metabolise arabinose (Δ*(araD-araB)567* genotype) [79], and therefore the inducer is not metabolised during the experiment.

First we tested the genetic interactions between *bipA* and all *E. coli* ABCFs in CFT073 background. At 37°C the *bipA* CFT073 knock-out strain has no growth defect (fig. S5A), however at 18°C, the Δ*bipA* strain displays a pronounced growth defect characteristic of strains defective in ribosome assembly (fig. 5A) [81]. Ectopic expression of Uup efficiently suppresses the growth defect, while deletion of *uup* in the *bipA* background exacerbates it (fig. 5A). Expression of EttA and Ybit have no effect, but expression of YheS leads to a dramatic growth defect. Importantly, in the wild type background, the expression of YheS has no effect on growth at 18°C (fig. S5B), indicating that the genetic interaction between *bipA* and *yheS* is specific. As reported previously, disruption of *bipA* leads to a dramatic ribosome assembly defect at low (18°C) temperature [71] (fig. 5B). The levels of mature 70S ribosomes as well as 50S subunits are dramatically decreased, accompanied by an accumulation of 50S assembly precursors (the peak marked with an asterisk) and free 30S subunits. Ectopic expression of Uup partially suppresses these defects, and in the Δ*bipA*Δ*uup* strain the defects are exacerbated. All of the effects described above are conditional on disruption of *bipA* since neither disruption of individual ABCF genes nor simultaneous disruption of *uup* and *ettA* – the only well-characterised ribosome-associated *E. coli* ABCF to date – causes cold-sensitivity (fig. S5C) or affects polysome profiles (fig. S5D).

**Fig. 5.**
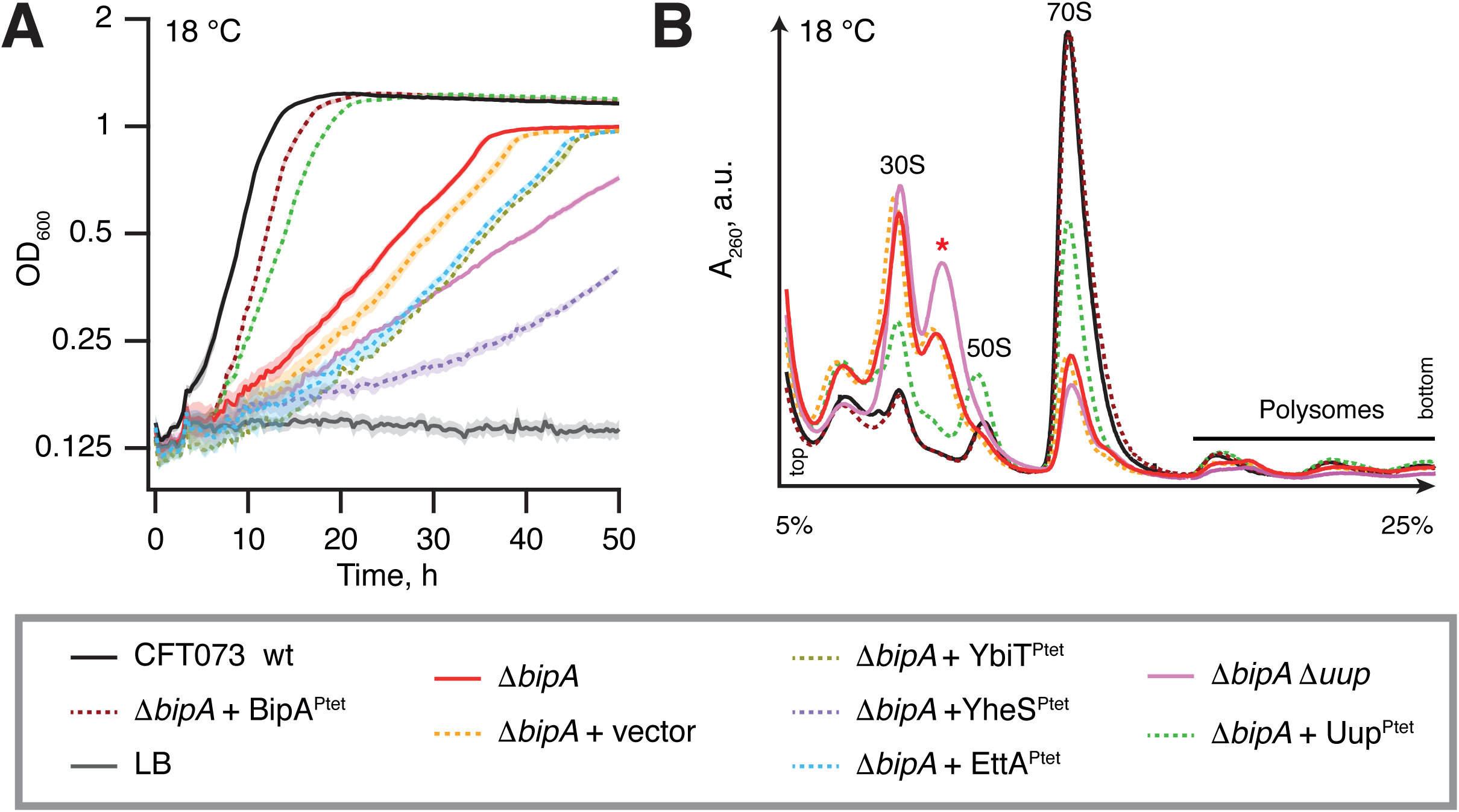
Overexpression of *E. coli* ABCF Uup suppresses cold sensitivity and ribosome assembly defects caused by loss of translational GTPase BipA. Growth (A) and sucrose gradient polysome analysis (B) of CFT073 wild type, isogenic Δ*bipA* and Δ*bipA*Δ*uup*, as well as CFT073 Δ*bipA* transformed with low-copy pSC vector expressing either BipA or ABCFs EttA, Uup, UheS and YbiT under control of constitutive promoter P_tet_. All experiments were performed in filtered LB at 18°C and data are presented as geometric means ± standard deviation (n = 3).

Next we set out to test the effects of EQ_2_ versions of ABCFs on translation. We validated the expression of the FTH-tagged ABCFs using Western blotting (fig. S6A). As was observed for untagged Uup (fig. 5A), the expression of FTH-tagged Uup suppresses the cold sensitivity caused by *bipA* deletion while YheS expression exacerbates the growth defect (fig. S6B-C). Expression of the EQ_2_ versions universally causes growth inhibition, both at 18°C in *ΔbipA* CFT073 (fig. S6C) and at 37°C in the wild type BW25113 background (fig. 6). Overexpression of none of the wild type ABCFs results in a growth defect (fig. 6A-D). Next we used a ^35^S-methionine pulse-labeling assay as a readout of translational inhibition. For all the ABCF-EQ_2_s, the methionine incorporation decreases, showing protein synthesis is clearly inhibited. The strongest effect is observed for EttA-EQ_2_ (fig. 6A) and YbiT-EQ_2_ (fig. 6C), and the weakest is seen in for Uup-EQ_2_ (fig. 6B).

**Fig. 6.**
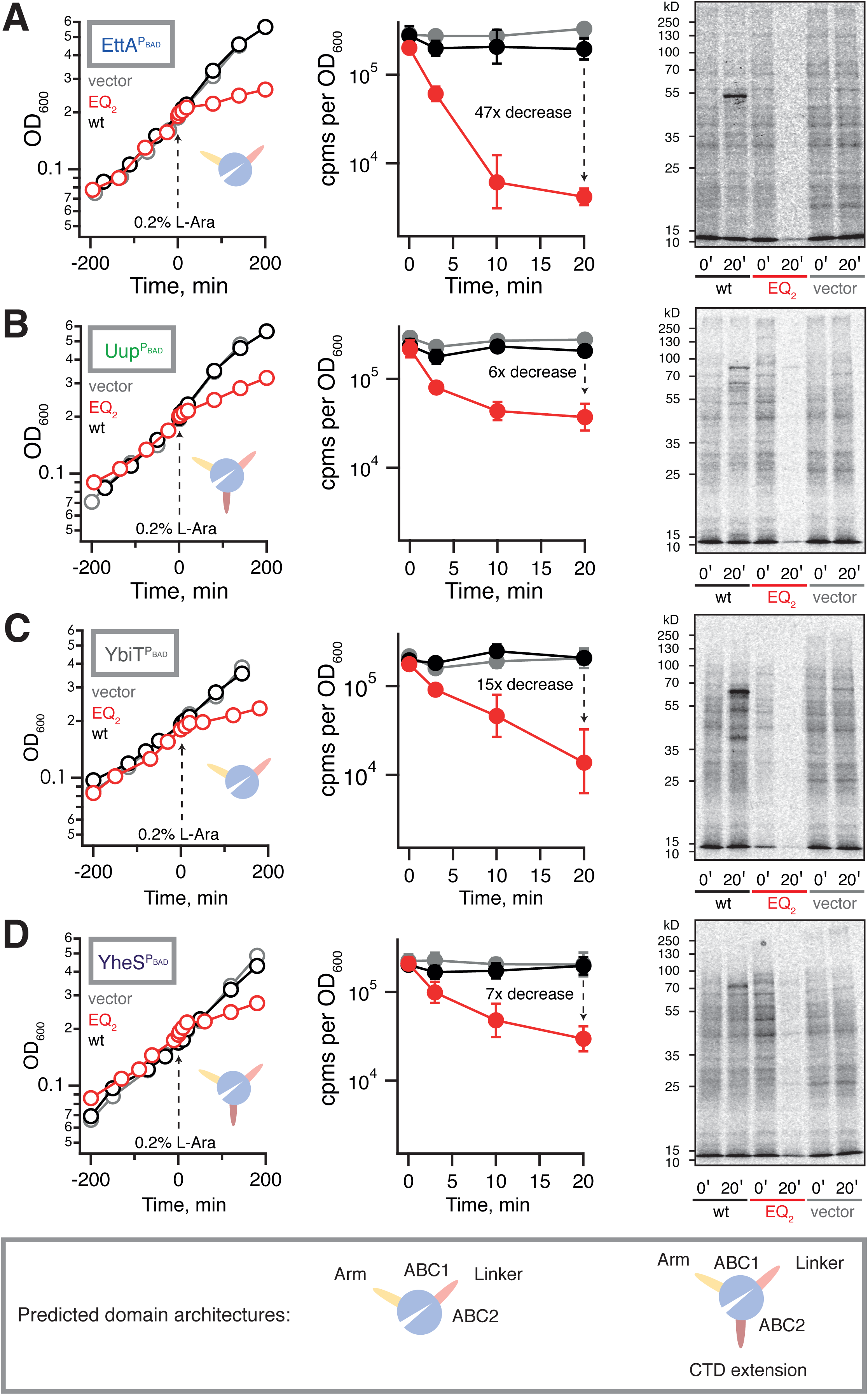
Expression of *E. coli* ABCF-EQ_2_ mutants inhibits growth and protein synthesis. Growth of wildtype *E. coli* BW2513 transformed with pBAD18 vector (grey trace) as well as *E. coli* BW2513 expressing either wild type (black trace) or EQ_2_ mutants (red trace) of EttA (A), Uup (B), YbiT (C), and YheS (D) under the control of arabinose-inducible promoter P_BAD_. Radiographs show the effect of wild type and EQ_2_ ABCF expression on protein synthesis, as probed by pulse labeling with L-[^35^S]-methionine. Expression was induced by the addition of L-arabinose to a final concentration of 0.2% at time point 0, and efficiency of incorporation was quantified by scintillation counting and visualised by autoradiography at 0 and 20-minute time points. Scintillation counting data are presented as geometric means ± standard deviation (n = 3). All experiments were performed at 37°C in Neidhardt MOPS medium [105] supplemented with 0.4% glycerol as a carbon source. The inset cartoons are a representation of ABCF domains and sub-domains, as per the legend in the lower box.

#### Phylogenetic analysis reveals putative AREs

Through phylogenetic analysis of predicted proteins and previously documented AREs with sequences available in UniProt [42] and the Comprehensive Antibiotic Resistance Database (CARD) [82], we have identified seven groups of AREs (fig. 1, table 1). Surprisingly, these can be quite variable in their subdomain architecture (fig. 7A). Some AREs (ARE1-5) have experienced extension of the Linker by on average around 30 amino acids compared to EttA, which is in line with the observation that the extended Linker is in close contact with the bound antibiotic in the case of ARE1 MsrE [33]. However, linker extension is not the rule for AREs; ARE7 (OptrA) has a linker of comparable length to the *E. coli* ABCFs (fig. 7A). This supports the notion based on the VmlR –ribosome co-structure that antibiotic protection by ABCFs can involve allosteric changes in the ribosome as well as through direct interaction [34]. The Arm subdomain that in EttA interacts with the L1 ribosomal protein and the L1 stalk rRNA (fig. 7A) varies in length among AREs, and the CTD extension may or not be present (fig. 7B). Surprisingly, the Uup protein from cave bacterium *Paenibacillus sp. LC231*, a sequence that is unremarkable among Uups, confers resistance to the pleuromutilin antibiotic tiamulin [39] (fig. S1).

**Fig. 7.**
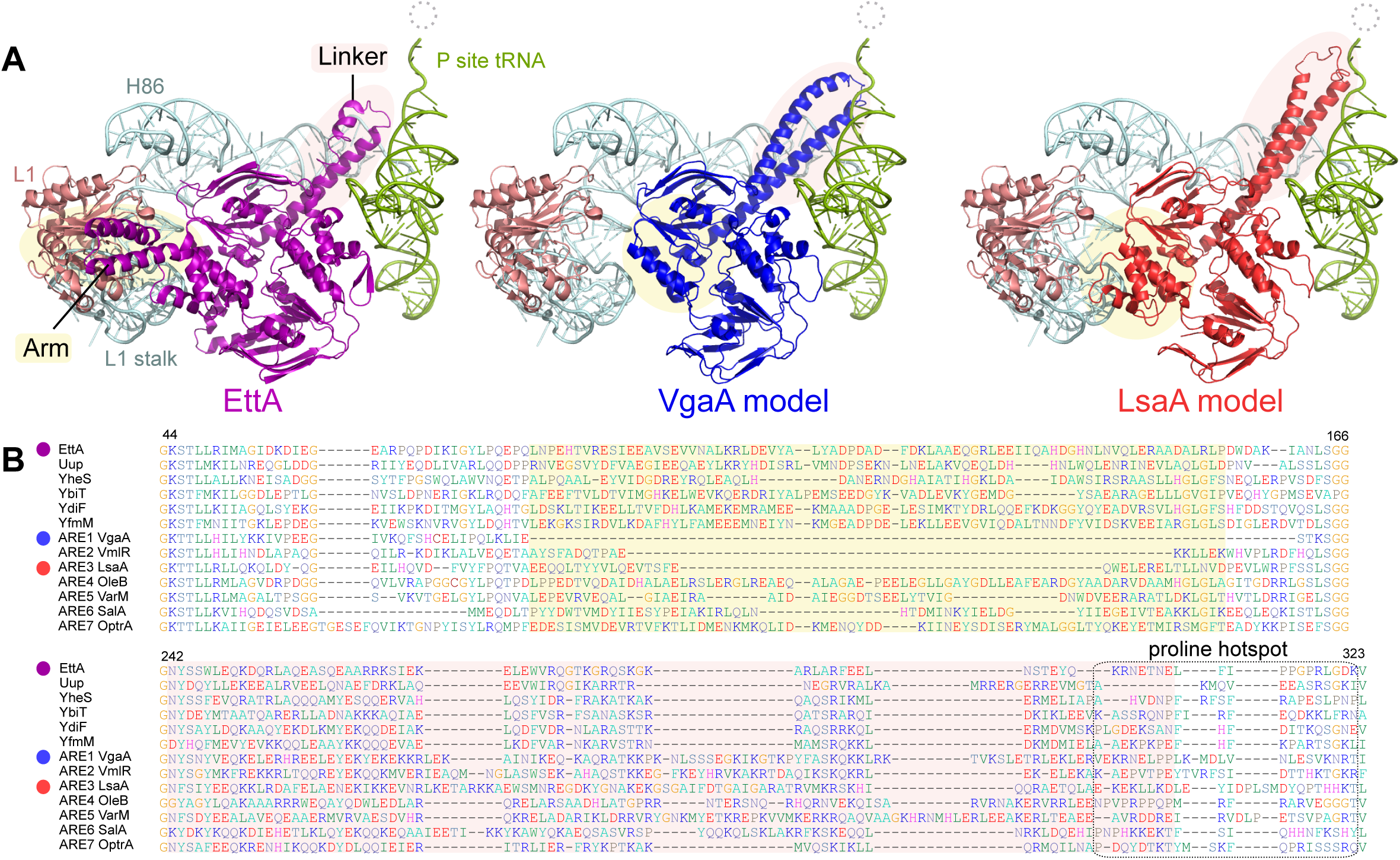
AREs tend to have relatively long linker regions that potentially extend towards the ribosome bound antibiotics. (A) The structure of EttA and its interacting ribosomal components from PDB 3J5S [17] is shown alongside homology models of *Staphylococcus aureus* VgaA and *Entercoccus faecalis* LsaA, using 3J5S as the template, with *de novo* modeling of the linker regions. The dotted circle shows the relative location of PTC-inhibiting antibiotics. Arm and linker regions are shaded in yellow and turquoise respectively. (B) Extracts from the multiple sequence alignment of *E. coli* and *B. subtilis* ABCFs, and representative AREs, containing the Arm (yellow shading) and Linker (turquoise shading) subdomains. Alignment numbering is according to the EttA sequence. A boxed region shows a region that is particularly rich in proline and polyproline in various ABCF family members.

Actinobacteria are the source of many ribosome-targeting antibiotics [83], and they have evolved measures to protect their own ribosomes, including ARE4 (OleB) and ARE5 (VarM). In addition to these known AREs, Actinobacteria encode a number of other ABCFs specific to this phylum (AAF1-6; table 1). It is possible that some – if not all – of these groups are in fact AREs. Other subfamilies may also be unidentified AREs, but two particularly strong candidates are BAF2 and BAF3 that have strong support for association with ARE5 (fig. 1) and are found in a wide range of bacteria (table 1). PAF1 is also worthy of investigation; it is found in the genomes of several pathogens in the proteobacterial genera *Vibrio, Enterobacter, Klebsiella, Serratia* and *Citrobacter*, but not *Escherichia* (table S1). With the exception of antibiotic producers, known antibiotic resistance ABCFs tend to have a variable presence within genera (fig. S7), probably because they are frequently transferred by mobile elements such as plasmids and transposons. Therefore, variability in the presence of a subfamily across species in a genus can be an indication that an ABCF is an ARE. Taking into account phylogenetic relationships (fig. 1, fig. S1) and disjunction within genera (fig. S7), we predict the following novel AREs: AAF1-5 (which tend to be found in antibiotic producers), BAF1-3, FAF1-2, and PAF1.

#### ABCFs are polyproline-rich proteins

Curiously, we find that ABCFs from both bacteria and eukaryotes are often rich in polyproline sequences, which are known to cause ribosome slow-down or stalling during translation [84]. This stalling is alleviated by the elongation factor EF-P in bacteria [85, 86], and indeed EF-P is required for full expression of EttA, which contains two XPPX motifs [87]. 56% of all the sequences in our ABCF database contain at least two consecutive prolines, compared to an overall 37% of all the proteins in the predicted proteomes considered here. There is a particularly proline-rich hotspot in the C-terminal part of the linker (fig. 7B), which can be up to nine consecutive prolines long in the case of Uup from *Novosphingobium aromaticivorans* (NCBI protein accession number WP_028641352.1). Arms and CTD extensions are also polyproline hotspots; PPP is a common motif in the Arm of EttA proteins, and polyprolines are frequent in YdiF, Uup and YheS CTD extensions. Prolines are rigid amino acids, and conceivably their presence may support the tertiary structures of ABCFs, particularly the orientation of the subdomain coiled coils [88].

## Discussion

### Towards a general model for non-eEF3 ABCF function

In the case of ABCFs that act on the assembled ribosome during translation, the E-site should be vacant (i.e. not filled by an E-site tRNA) for the protein to bind. Specifically, this would be when the E-site has not yet received a tRNA (during initiation, where ABCF50 (ABCF1) functions [13]) or when an empty tRNA has dissociated and not been replaced (during slow or stalled translation such as in the presence of an antibiotic, as in the case of AREs and EttA [16, 17, 34]). EttA has been proposed to promote the first peptide bond after initiation through modulation of the PTC conformation [16, 17]; similarly, allosteric effects acting on the PTC have been observed for the ARE VmlR [34]. This structural modulation or stabilisation could conceivably be a general function of ABCFs, with the specific ribosomal substrate differing depending on the stage of translation, assembly, or cellular conditions. Differences in subdomains would determine both what is sensed and the resulting signal. For instance the presence of the Arm would affect signal transmission between the PTC and the L1 protein and/or the L1 stalk.

### Evolution of AREs

In order to determine antibiotic resistance capabilities and track the transfer routes of resistance, it is critical to be able to annotate antibiotic resistance genes in genomes. This requires discrimination of antibiotic genes from homologous genes from which resistance functions have evolved. At present this is not straightforward for ABCF AREs, as the distinction between potential translation and antibiotic resistance factors is ambiguous. ABCF AREs have been compared to the Tet family of antibiotic resistance proteins that evolved from trGTPase EF-G to remove tetracycline from the ribosome [29]. However the distinction between Tet proteins and EF-G is much more clear-cut, with Tet comprising a distinct lineage in the evolutionary history of trGTPases [35]. The surprising lack of a clear sequence signature for antibiotic resistance in the ARE ABCFs suggests that antibiotic resistance functions may evolve in multiple ways in ABCFs, and that ABCFs closely related to AREs may have similar functions to the AREs, while not conferring resistance. For example, there are multiple small molecules that bind the PTC and exit tunnel [89], and conceivably ABCFs may be involved in sensing such cases, removing the small molecule, or allowing translation of a subset of mRNAs to continue in its presence. Even macrolide antibiotics that target the exit tunnel do not abrogate protein synthesis entirely, but rather reshape the translational landscape [90, 91].

## Conclusion

ABCFs are stepping into the limelight as important translation, ribosome assembly and antibiotic resistance factors. We have found hydrolysis-incompetent EQ_2_ mutants of all four *E. coli* ABCFs inhibit protein synthesis, suggesting they all function on the ribosome. Overexpression of Uup suppresses both the cold-sensitivity and the 50S ribosome assembly defect caused the loss of translational GTPase BipA, suggesting that Uup is involved in the 50S ribosome subunit assembly, either directly or indirectly, for example by fine-tuning expression of ribosomal proteins. Additionally, we have added ARE2 VmlR to the repertoire of AREs confirmed to act on the ribosome, joining the ranks of LsaA, VgaA and MsrE, all ARE1s. Considering the well-established ribosome association of eukaryotic ABCFs, our results suggest that ribosome-binding is a general - perhaps ancestral - feature of ABCFs. However, the ABCF family is diverse, and even within subfamilies, there can be differences in subdomain architecture. We have identified clusters of antibiotic resistance ARE ABCFs, and predicted likely new AREs. Strikingly, the AREs do not form a clear monophyletic group, and either antibiotic resistance has evolved multiple times independently from the ABCF diversity, or this is an innate ability of ABCFs, raising the possibility of a general role of ABCFs in ribosome-binding small molecule sensing and signaling.

## Methods

### Sequence searching and classification

Predicted proteomes were downloaded from the NCBI genome FTP site (2^nd^ December 2014). One representative was downloaded per species of bacteria (i.e. not every strain). Seven additional proteomes were downloaded from JGI (*Aplanochytrium kerguelense, Aurantiochytrium limacinum, Fragilariopsis cylindrus, Phytophthora capsici, Phytophthora cinnamomi, Pseudo-nitzschia multiseries,* and *Schizochytrium aggregatum*). Previously documented AREs were retrieved from UniProt [42] and the Comprehensive Antibiotic Resistance Database (CARD) [82]. Taxonomy was retrieved from NCBI, and curated manually where some ranks were not available.

An initial local BlastP search was carried out locally with BLAST+ v 2.2.3 [92] against a proteome database limited by taxonomy to one representative per class or (order if there was no information on class from the NCBI taxonomy database), using EttA as the query. Subsequent sequence searching against the proteome collections used hmmsearch from HMMER 3.1b1, with HMMs made from multiple sequence alignments of subfamilies, as identified below. The E value threshold for hmmsearch was set to 1e_-70_, a value at which the subfamily models hit outside of the eukaryotic-like or bacterial-like ABCF bipartitions, ensuring complete coverage while not picking up sequences outside of the ABCF family.

Sequences were aligned with MAFFT v7.164b (default settings) and Maximum Likelihood phylogenetic analyses were carried out with RAxML-HPC v.8 [93] on the CIPRES Science Gateway v3 [94] using the LG model of substitution, after removing positions containing >50% gaps. Additional phylogenetic analyses of representative sequences were carried out as described in the section “phylogenetic analysis of representatives”, below. Trees were visualised with FigTree v. 1.4.2 (http://tree.bio.ed.ac.uk/software/figtree/) and visually inspected to identify putative subfamilies that preferably satisfied the criteria of 1) containing mostly orthologues, and 2) had at least moderate (>60% bootstrap support). These subfamilies were then isolated, aligned separately and used to make HMM models. Models were refined with subsequent rounds of searching and classification into subfamilies first by comparisons of the E values of HMM hits, then by curating with phylogenetic analysis. After the final classification of all ABCF types in all predicted protein sequences using HMMER, some manual correction was still required. For example, cyanobacterial sequences always hit the chloroplast HMM with a more significant E value than the bacterial model, and eEF3/New1/eEF3L sequences could not be reliably discriminated between using E value comparisons. Therefore, the final classification is a manually curated version of that generated from automatic predictions (table S2). All sequence handling was carried out with bespoke Python scripts, and data was stored in a MySQL database (exported to Excel files for the supplementary material tables). Identification of EFL and eEF1A in predicted proteomes was carried out with HMMER, using the HMMs previously published for these trGTPases [35], The E value cut-of was set to e-200, lenient enough to match both eEF1A or EFL, with assignment to either protein subfamily made by E value comparisons as above.

### Domain prediction

Domain HMMs were made from subalignments extracted from subfamily alignments. Partial and poorly aligned sequences were excluded from the alignments. All significant domain hits (<E value 1e_-3_) for each ABCF sequence were stored in the MySQL database. Sequence logos of domains were created with Skylign [95]. Putative transit peptides for mitochondrial and plastid subcellular localization were predicted with the TargetP web server hosted at the Technical University of Denmark [96].

### Phylogenetic analysis of representatives

For the representative tree of the ABCF family, taxa were selected from the ABCF database to sample broadly across the tree of life, including eukaryotic protistan phyla, while also covering all subfamilies of ABCFs. Sequences were aligned with MAFFT with the L-ins-i strategy [97] and positions with >50% gaps, and several ambiguously aligned positions at the termini were removed. The resulting 249 sequences, and 533 positions were subject to RAxML and IQ-TREE Maximum Likelihood phylogenetic analysis, both run on the CIPRES Science Gateway v3 [94]. RAxML was run with the LG substitution matrix, as favoured by ProtTest 2.4 [98] and 100 bootstrap replicates. Bootstrapping with RaxML yields a value (maximum likelihood bootstrap percentage, MLB) for how much of the input alignment supports a particular branch in the tree topology, and therefore the reliability of that branch. These support values are indicated on branches in the tree figures. In the case of IQ-TREE, the most appropriate model was selected by the program during the run, which also favoured the LG substitution matrix. IQ-TREE was run with its ultrafast bootstrapping approximation method to ascertain support values (UFB) for branches out of 1000 replicates [99]. To test whether our trees made with ABC domains separately are incompatible (as might indicate recombination), the RaxML and IQ-TREE analyses were repeated with a dataset containing the ABC domains uncoupled from each other and aligned together. Alignments were prepared as above, to make a data set of 204 alignment positions from 525 taxa (Text S1).

Bayesian inference phylogenetic analysis was carried out with MrBayes v3.2.6, also on the CIPRES gateway. The analysis was run for 1 million generations, after which the standard deviation of split frequencies (SDSF) was 0.08. The mixed model setting was used for determining the amino acid substitution matrix, which converged on WAG. RAxML analysis with the WAG model showed no difference in topology for well-supported branches compared to the RAxML tree with the LG model. Branch support values in this case are posterior probabilities, shown on the tree figure as Bayesian Inference posterior probabilities (BIPP), on a scale of 0 to 1, with increasing probability.

For the rooted tree with ABCE as the outgroup, all ABCFs were selected from *Arabidopsis thaliana, Homo sapiens, Escherichia coli, Bacillus subtilis, Saccharomyces cerevisae, Schizosaccharomyces pombe* and *Methanococcus maripaludis*. Ambiguously aligned sites were identified and removed manually. RAxML, IQ-TREE and MrBayes were carried out as above on the resulting 344 positions from 35 sequences. The MrBayes analysis stopped automatically when the SDSF dropped to the 0.009 threshold, which was at 235,000 generations.

For the tree of eukaryotic and viral ABCFs rooted with YheS, sequences were extracted from the ABCF database, and viral sequences were found in the NCBI protein database using BlastP. The resulting 658 sequences were aligned with MAFFT and a RAxML analysis of 645 positions was carried out as above. The alignments used to build the phylogenies presented in the main text as supplementary information are available in text S1, along with trees that are used for ascertaining branch support but not included as figures.

### Structural analyses

Homology modeling was carried out using Swiss Model [100] with EttA (PDB ID 3J5S) as the template structure. Because the linkers were lacking secondary structure, QUARK [101] was used for *ab initio* structure modeling of these regions, and the resulting coils were aligned back to the homology model using the structural alignment method of MacPyMOL [102]. The presence of coiled coil regions was predicted with the COILS program hosted at the ExPASy Bioinformatics Research Portal [103].

### Construction of plasmids and bacterial strains

All bacterial strains and plasmids used in this study are described in **Supplementary Methods** and listed in **Table 2**.

**Table 2.**
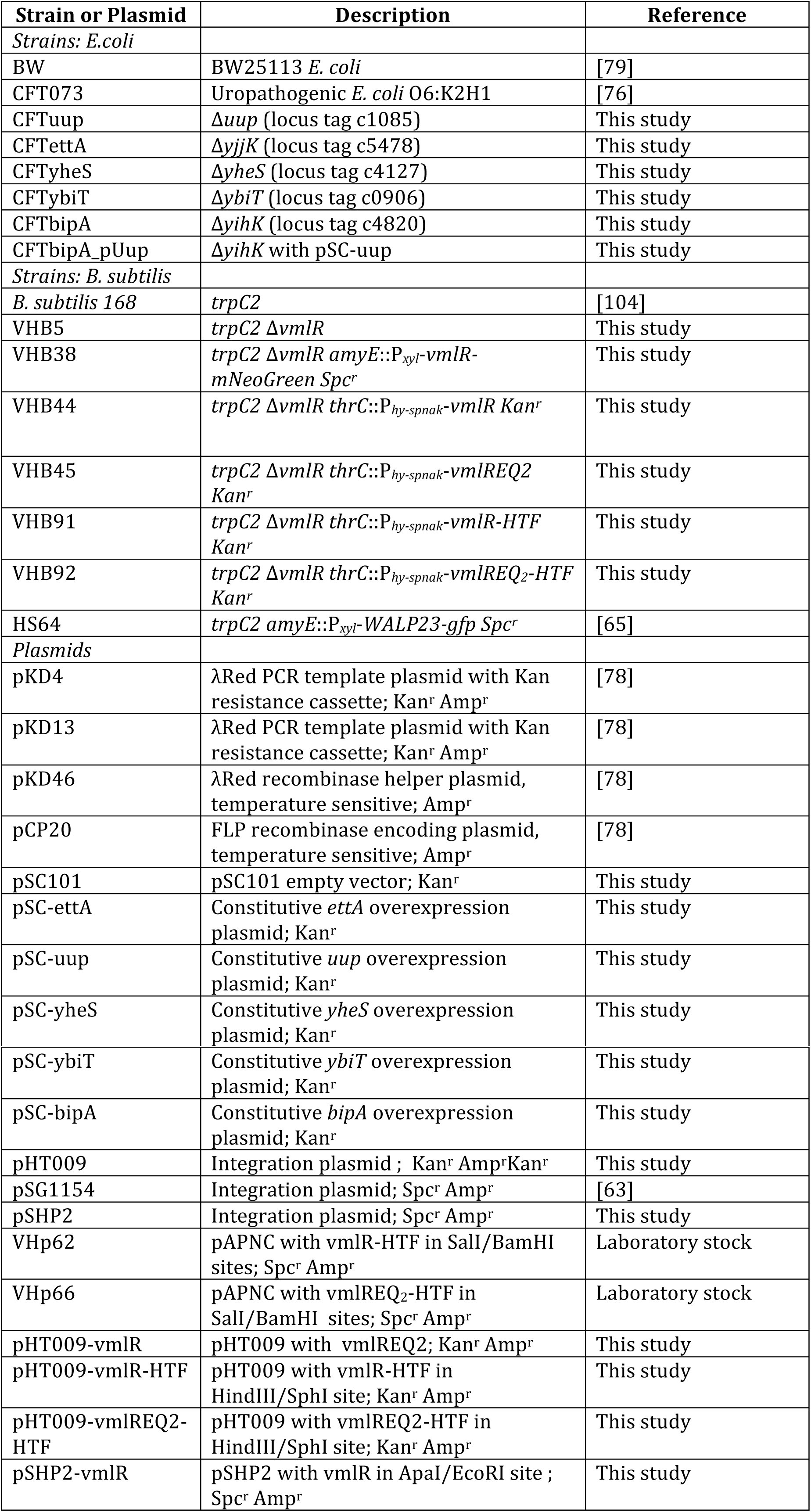
Strains and plasmids used in the study.

### Growth assays

Bacterial growth (OD_600_) was monitored using a Bioscreen C (Oy Growth Curves Ab Ltd) microplate reader in Honeycomb plates (150 µL culture per well) with continuous shaking (speed: fast, amplitude: normal). All experiments with CFT073 were performed at 18°C unless stated otherwise. Three biological replicates were averaged for each growth curve and the data presented as geometric means ± standard deviation.

#### E. coli *transformed with pSC101-based expression plasmids*

Overnight (16 h) cultures were pre-grown in LB medium supplemented with 50 μg/mL kanamycin, diluted to OD_600_ of 0.03 in filtered LB and grown in Bioscreen C microplate reader as described above.

#### E. coli *transformed with pBAD-based expression plasmids*

Overnight (16 h) cultures were pre-grown in Neidhardt MOPS medium [105] supplemented with 0.1% of casamino acids, 0.4% glucose as a carbon source and 100 μg/mL carbenicillin, diluted to OD_600_ of 0.03 in the same media but containing and 0.5% glycerol instead of 0.4% glucose as well as supplemented with 0.5% arabinose and grown in Bioscreen C microplate reader as described above.

#### *Antibiotic resistance testing of tagged* B. subtilis *VmlR*

*B. subtilis* strains VHB38, VHB91 and VHB92 were pre-grown on LB plates overnight at 30°C. Fresh individual colonies were used to inoculate filtered LB medium, either in the presence and absence of 1 mM IPTG (for VHB91 and VHB92) or in the presence and absence of 0.3% xylose (for VHB38), and OD_600_ adjusted to 0.01. The cultures were seeded on Honeycomb plates, and plates incubated in a Bioscreen C at 37 °C with continuous shaking as described above for *E. coli* cultures. After 90 minute incubation (OD_600_ ≈ 0.1) increasing concentrations of lincomycin (final concentration 0 - 5 μg/ml) were added and growth was monitored for additional 6 hours. Three biological replicates were averaged for each growth curve and the data presented as geometric means ± standard deviation.

### Fluorescence microscopy

Fluorescence microscopy was carried out with cell grown to early-mid logarithmic growth phase (OD_600_ of 0.2-0.5) in LB medium at 37 °C in the presence or absence of inducers. The used inducer concentrations were 0.3% for VmlR-mNG and 1% for WALP23-GFP. If indicated, the cells were incubated with 5µg/ml lincomycin upon shaking at 37 °C prior to the microscopy. The cells were immobilised on microscopy slides covered with a thin film of 1.2% (w/v) agarose in H_2_O as described in detail elsewhere [106]. The microscopy was carried out with Nikon Eclipse Ti equipped with Nikon Plan Apo 100x/1.40 Oil Ph3 objective, Sutter Instrument Company Lambda LS xenon arc light source, and Photometrics Prime sCMOS camera. The images were captured using Metamorph 7.7 (Molecular Devices) and analysed using Fiji [107].

### Western blot analysis of FTH-tagged wt and EQ_2_ E. coli ABCF proteins

#### Preparation of bacterial samples

Bacteria were grown either at 37 °C (*E. coli* BW25113 derivatives) or 18°C (*E. coli ΔbipA* CFT073 derivatives) up to OD_600_ of 0.5 in 50 mL of Neidhardt MOPS minimal medium [105] supplemented with 0.1% casamino acids (w/v), 0.5% glycerol (w/v) and 100 µg/mL carbenicillin and L-arabinose was added to a final concentration of 0.2% (w/v). Cultures grown at 37°C were harvested 10 minutes after induction by pouring them into precooled centrifuge bottles containing 100 g of crushed ice and centrifuged at 10,000 rpm for 10 minutes at 4 °C (Beckman JLA16.250 rotor). Cultures grown at 18°C were harvested 5 hours after induction by collecting into precooled centrifuge bottles and pelleting at 10,000 rpm for 10 minutes at 4 °C (Beckman JA25.50 rotor). Lysates were prepared the same as for polysome profiling of *E. coli* (see below).

#### Western blotting

3 μg of total protein as determined by Bradford assay of each sample was resolved on 10% SDS-PAGE gel and transferred to 0.2 μm nitrocellulose membrane (Trans-Blot^®^ Turbo™ Transfer Pack, Bio-Rad) using Turbo MIXED MW protocol in Trans-Blot^®^ Turbo™ Transfer System (Bio-Rad). The membrane was blocked in PBS-T (1x PBS 0.05% Tween-20) with 5% w/v nonfat dry milk at room temperature for one hour. Antibody incubations were performed for one hour in 1% nonfat dry milk in PBS-T with five 5-minute washes in fresh PBS-T between and after antibody incubations. FTH-tagged ABCFs were detected using anti-Flag M2 primary (Sigma-Aldrich, F1804; 1:10,000 dilution) antibodies combined with anti-mouse-HRP secondary (Rockland; 610-103-040; 1:10,000 dilution) antibodies. AECL detection was performed on ImageQuant LAS 4000 (GE Healthcare) imaging system using Pierce^®^ ECL Western blotting substrate (Thermo Scientific).

### Polysome profiling analysis of E. coli strains

#### Preparation of bacterial samples

Overnight (16 h) cultures were pre-grown at 37°C in LB medium supplemented with 50 μg/mL kanamycin in the case of strains transformed with pSC101-based expression plasmids. Overnight cultures were diluted in filtered LB (33 mL cultures) and after 24 h growth at 18 °C harvested by pouring into precooled centrifuge bottles and pelleting at 10,000 rpm for 10 minutes at 4 °C (Beckman JA25.50 rotor). For the sake of convenience, cultures were diluted to different starting densities (Table S3) to ensure that all of them reach OD_600_ ≈0.5 simultaneously.

#### Preparation of clarified lysates

Cell pellets were resuspended in 0.4 mL of Polymix buffer [108] (20 mM HEPES:KOH pH 7.5, 95 mM KCl, 5 mM NH_4_Cl, 5 mM Mg(OAc)_2_, 0.5 mM CaCl_2_, 8 mM putrescine, 1 mM spermidine, 1 mM DTT) and 200 µL of pre-chilled zirconium beads (0.1 mm) were added to each sample. Cellular lysates were prepared by a FastPrep homogeniser (MP Biomedicals) (three 20 seconds pulses at speed 6.0 mp/sec with chilling on ice for 1 minutes between the cycles) and clarified by centrifugation at 21,000 g for 10 minutes at 4 °C. The supernatant was carefully collected avoiding the lipid layer and cellular pellet, aliquoted, frozen in liquid nitrogen and stored at -80 °C until further processing.

#### Sucrose gradient centrifugation

After melting the frozen samples on ice, 2 A_260_ units of each extract was loaded onto 5–25% (w/v) sucrose density gradients in Polymix buffer, 5 mM Mg_2+_ [108]. Gradients were resolved at 35,000 rpm for 2.5 hours at 4 °C in SW41 rotor (Beckman) and analysed using Biocomp Gradient Station (BioComp Instruments) with A_260_ as a readout. The ribosome profiles presented were normalised to the total area under the curve and are representative of at least three independent experiments for each strain.

### Polysome profiling and Western blot analysis of B. subtilis strains

Experiments were performed as described above for *E. coli* strains, with minor modifications.

#### Preparation of bacterial samples and preparation of clarified lysates

VHB90 and VHB91 strains were pre-grown on LB plates overnight at 30 °C. Fresh individual colonies were used to inoculate 200 mL LB cultures. The cultures were grown at 37 °C until OD_600_ of 0.3 and IPTG was added to final concentration of 30 μM. After 30 min cells were collected by centrifugation (8,000 rmp, 10 minutes), dissolved in 0.5 mL of Polymix buffer [108] (20 mM HEPES:KOH pH 7.5, 95 mM KCl, 5 mM NH_4_Cl, 10 mM Mg(OAc)_2_, 0.5 mM CaCl_2_, 8 mM putrescine, 1 mM spermidine, 1 mM DTT, 2 mM PMSF), lysed (FastPrep homogeniser (MP Biomedicals): four 20 seconds pulses at speed 6.0 mp/sec with chilling on ice for 1 minutes between the cycles), and clarified by ultracentrifugation (14,800 rpm, 20 minutes).

#### Sucrose gradient centrifugation and Western blotting

Clarified cell lysates were loaded onto 7–35% sucrose gradients in Polymix buffer [108] (20 mM HEPES:KOH pH 7.5, 95 mM KCl, 5 mM NH_4_Cl, 10 mM Mg(OAc)_2_, 0.5 mM CaCl_2_, 8 mM putrescine, 1 mM spermidine, 1 mM DTT) and subjected to centrifugation (35,000 rpm for 3 hours at 4 °C). C-terminally HTF-tagged VmlR (wild type and EQ_2_ mutant) and ribosomal protein L3 of the 50S ribosomal subunit were detected using either anti-Flag M2 primary combined with anti-mouse-HRP secondary antibodies or anti-L3 primary (a gift from Fujio Kawamura) combined with goat anti-rabbit IgG-HRP secondary antibodies, respectively. All antibodies were used at 1:10,000 dilution.

### L-[^35^S]-methionine pulse-labelling

#### Preparation of bacterial samples

Since glucose specifically inhibits the arabinose promoter, the cultures were grown in defined Neidhardt MOPS medium [105] supplemented with 0.4% glycerol as a carbon source. One colony of freshly transformed *E. coli* BW25113 cells expressing N-terminal FTH-tagged ABCFs (wild type and EQ_2_ mutants) from pBad vector was used to inoculate 10 mL of 1x MOPS media supplemented with 0.4% glycerol 100 μg/mL carbenicillin, and the cultures were grown until early stationary phase (about 24 hours). Stationary phase cells were diluted to OD_600_ of 0.04–0.07 in 25 mL of the same media, grown at 37 °C with vigorous shaking (200 rpm) to OD_600_ of 0.15–0.2 and expression of ABCFs was induced by addition of L-arabinose to the final concentration of 0.2%.

#### L-[^35^S]-methionine pulse-labelling

For radioactive pulse labelling, 1 μCi L-[^35^S]-methionine (500μCi, PerkinElmer) aliquots were prepared in 1.5 mL Eppendorf tubes. As a zero time point, 1 mL of cell culture was taken and mixed with an aliquot of radioactive methionine just before inducing cells with L-arabinose. Simultaneously a 1 mL aliquot was taken for an OD_600_ measurement. All consecutive samples were processed similarly at designated time points after induction. ^35^S-methionine incorporation was stopped after 5 minutes by chloramphenicol added to the final concentration of 200 µg/mL. Subsequent processing of samples differs in the case of scintillation counting and autoradiography.

#### Scintillation counting

1 mL of culture was combined with 200 µL of 50% trichloroacetic acid (TCA), passed through a GF/C filter (Whatman) prewashed with 5% TCA and unincorporated label was removed by washing the filter with 5 mL of ice-cold 5% TCA followed by 5 mL of ice-cold 95% EtOH [109]. Filters were dried for at least 2 hours, and counted on a TRI-CARB 4910TR 110 V scintillation counter (PerkinElmer) (5 mL of ScintiSafe 3 scintillation cocktail (FisherScientific) per sample, pre-soaked with shaking for 15 minutes prior to counting).

#### Autoradiography

1 mL cultures were pelleted by centrifugation, cell pellet washed with Phosphate Buffered Saline (PBS) to remove unincorporated L-[^35^S]-methionine and dissolved/lysed in 50 μL 1x SDS-loading buffer. Samples were normalised by OD600 by addition of appropriate volume of 1x SDS-loading buffer (50-80 µL according to OD600), and 10 µL of the sample was loaded onto 10% SDS-PAGE and resolved electrophoretically (BioRad), gels were dried on Whatman paper, exposed on BAS storage phosphor screen (GE Heathcare) overnight, and scanned by Typhoon imaging system (GE Heathcare).

## Acknowledgements

We are grateful to the anonymous reviewers in their suggestions of additional experiments and analyses that strengthened the manuscript. We gratefully acknowledge Fujio Kawamura for providing anti-L3 primary antibodies and Stijn Hendrik Peeters for constructing the pSHP2 plasmid. Thanks to Martin Carr for discussion and advice about eEF3 and EFL co-distribution.

## Author contributions

GCA conceived the study and carried out all the bioinformatic analyses except for sequence logo construction and eEF1A/EFL/eEF3 co-distribution analysis, which was carried out by CKS. VH, VM, MK, HS, HT, TT and MP designed experimental analyses. VM, MK, HT, MH, TS, MR, HS and JWG carried out the experiments. VH, GCA, HS, TT, HT, VM and MK analyzed the data. GCA and VH drafted the manuscript with input from VM, MK, HT, MH, CKS, JWG, TS, MP, TT and HS.

## Supplementary file captions

**S1 Table. Taxonomy of 4505 species, and their ABCF composition**

All species considered in the analysis are listed, ordered by taxonomy. The number and identity of ABCF subfamilies are recorded.

**S2 Table. Classification of 16848 ABCF sequences into subfamilies, accompanied by domain assignments**

The unique sequence identifiers are included for retrieval of data from online repositories.

**S3 Table. Domain coordinates**

The domain coordinates for the representative sequences in fig. 3A are listed. In the second tab, all the coordinates for each identified domain in all ABCFs are given. As these are from HMM hits, the same domain can have more than one hit in each protein. For example, the ABC1 HMM always hits the ABC2 domain, and vice versa. Duplicate domain hits were removed when generating fig 3A.

**S4 Table. Presence and absence of EFL, eEF1A and eEF3 in eukaryotes**

Where the distribution is unchanged within a specific taxonomic lineage, those rows are collapsed down to one, and the highest common taxonomic rank is given. The full lineage data is available in the second tab.

**S5 Table. Transit peptide predictions** Predictions were made separately for plastid-containing and non-plastid containing eukaryotes. The description of the output format is shown below the predictions.

**Table S6. Primers used in the study**

**Table S7. The starting OD_600_ of *E. coli* CFT073 and its derivatives.**

**S1 Figure. Ladderised version of the Figure 1 tree**

All branch support values and taxon names including subfamily identity are shown. Branch colouring is as per Figure 1. Orange stars show Uup sequences that have truncated Arm subdomains.

**S2 Figure. Sequence logos of domains show amino acid biases**

Sequence logos of each domain HMM. The height of stacked amino acids at each position show the information content, in bits, with letters dividing the height according to their estimated probability. Beneath the stacked amino acids there are three lines showing probabilities; line 1 is occupancy, the probability of observing a letter –rather than a gap -at that position. Line 2 is the probably of seeing an insertion at that position, and line 3 is the expected length of an insertion following that position.

**S4 Figure. Functional testing of inducible *B. subtilis* VmlR-HTF and VmlR-NeonGreen fusion proteins.** All experiments were performed in filtered LB at 37°C in the presence of increasing concentrations of lincomycin and presented as the geometric mean ± standard deviation (n = 3). HTF stands for C-terminal His_6_-TEV-3xFLAG tag.

**S5 Figure. Growth and polysome analysis of *E. coli* CFT073 wild type, Δ*bipA*, Δ*abcf* and Δ*bipA*Δ*uup* strains, and *E. coli* CFT073 wild type overexpressing ABCFs under the control of constitutive P_tet_ promoter.** All experiments were performed in filtered LB at either 18°C (A) or 37°C (B-D) and growth data are presented as geometric means ± standard deviation (n = 3).

**S6 Figure. Functional testing of FTH-tagged *E. coli* ABCF wild type and EQ_2_ proteins.**

Western blot (A) and growth assays (B and C) N-terminally FTH-tagged *E. coli* ABCF proteins expressed under the control of arabinose-inducible P_BAD_ promoter. Growth experiments were performed in MOPS media at 18°C and presented as geometric means ± standard deviation (n = 3). Western blotting was performed either at 18°C or 37°C. FTH stands for N-terminal 3xFLAG-TEV-His_6_ tag.

**S7 Figure. AREs show variability in presence across species of the same genus, which can be used to predict novel ARE ABCFs.** Genera were selected that contained more than 20 species in the ABCF database. For each subfamily in each of those genera, the percent of species in which that subfamily is found, is plotted. Colder colours are those ABCFs that are more universal, as typical of housekeeping genes. AREs on the other hand have a more patchy distribution, as shown by their tendencies for warmer colours. Antibiotic producing genera are the exception, where AREs and ARE-like subfamilies can be universal within a genus.

**S3 Figure. Maximum likelihood phylogeny of eukaryotic subgroup-type ABCFs from eukaryotes and viruses.**

The tree is rooted with bacterial YheS sequences. Branch support is from 100 bootstrap replicates and branch length is proportional to the number of amino acid substitutions (see lower scale bar). Names are coloured by subfamily.

**S1 Text. Supplementary sequence alignments and phylogenetic trees.**

The file contains sequence alignments used to generate Figures 1 (and S1), 2 and S3 in FASTA format, and Newick format phylogenetic trees that are not shown as figures. Each entity (alignment or tree) is separated by comments preceded by “#”.

**S1 Materials and Methods**

**Supplementary methods describing the construction of plasmids and bacterial strains**

